# Systematic Analysis of Metabolic Pathway Distributions of Bacterial Energy Reserves

**DOI:** 10.1101/545715

**Authors:** Liang Wang, Jianye Yang, Yue Huang, Qinghua Liu, Yaping Xu, Xue Piao, Michael J. Wise

## Abstract

Metabolism of energy reserves is essential for bacterial functions, such as pathogenicity, metabolic adaptation, and environmental persistence, *etc*. Previous bioinformatics studies have linked gain or loss of energy reserves such as glycogen and polyphosphate (polyP) with host-pathogen interactions and bacterial virulence based on a comparatively small number of bacterial genomes or proteomes. Thus, understanding the theoretical distribution patterns of energy reserves across bacterial species provides a shortcut route to look into bacterial lifestyle and physiology. So far, five major energy reserves have been identified in bacteria due to their capacity to support bacterial persistence under nutrient deprivation conditions. These include polyphosphate (polyP), glycogen, wax ester (WE), triacylglycerol (TAG), and polyhydroxyalkanoates (PHAs). Although the enzymes related with metabolism of energy reserves are well understood, there is a lack of systematic investigations into the distribution of bacterial energy reserves from an evolutionary point of view. In this study, we sourced 8282 manually reviewed bacterial reference proteomes from UniProt database and combined a set of hidden Markov sequence models to search homologs of key enzymes related with the metabolism of energy reserves. The distribution patterns were visualized in taxonomy-based phylogenetic trees. This study reveals that specific pathways and enzymes are restricted within certain types of bacterial groups, which provides evolutionary insights into the understanding of their origins and functions. In addition, the study also confirms that loss of energy reserves is correlated with bacterial genome reduction. Through this analysis, a much clearer picture about the metabolism of energy reserves in bacteria is presented, which could serve as a guide for further theoretical and experimental analyses of bacterial energy metabolism.

## Introduction

Due to the diversity of environmental niches that bacteria have colonized through millions of years of adaptation and evolution, bacteria have evolved specialized sets of metabolic pathways to live optimally in these environments, which can be reflected in their characteristic genomes, gene transcription profiles, and also proteomes [1]. Previously, comparative genomics studies have shown that loss of glycogen metabolism could serve as an indicator for bacterial parasitic lifestyle while gain of polyphosphate (polyP) metabolism seems to link with free-living lifestyle and higher bacterial virulence [2, 3]. In addition, it has been observed that loss of glycogen or polyP metabolism is associated with reduced genome size, providing a hint about genome reduction [2, 4]. It is well known that both glycogen and polyP are important energy sources in bacteria. Thus, presence or absence of energy reserves in bacteria could be important for *in silico* analysis of bacterial physiology and lifestyle, especially when large numbers of sequenced bacterial genomes are available and many of those are unculturable by traditional laboratory techniques.

It is well known now that energy reserves play essential roles in bacteria for their regular activities to sense and respond to changing environments and different types of stresses, such as temperature fluctuation and nutrient deprivation, *etc.* [4]. Although there are many different energy-related compounds, not all of them can be classified as energy reserves. According to Wilkinson, three principles should be satisfied for a compound to be considered as an energy reserve, which are: 1) accumulation when energy is over-supplied, 2) utilization when energy is insufficient, and 3) apparent advantages by consuming the compound when compared with those without it [5]. Through physiological and biochemical studies, five energy storage compounds have been regarded to meet the criteria, which are wax ester (WE), triacylglycerol (TAG), polyhydroxyalkanoates (PHAs), polyphosphates (polyP) and glycogen [2, 4-6].

Several studies have investigated the distribution patterns of energy reserves in bacteria, most of which were based on small sets of bacterial genomes or proteomes. No systematic analysis from the evolutionary point of view currently exists [2, 4, 7-10]. In this study, we collected 8282 manually reviewed bacterial proteomes from UniProt database and sourced key enzymes directly involved in the metabolism of five energy reserves from public literature and database (**Table 1**) [11]. Hidden Markov sequence models were used for searching homologs in bacterial proteomes. Distribution patterns of representative metabolic pathways of the fiver major reserves are presented in **Supplementary Table S1**. In order to gain an explicit view about distributions of the enzymes along the evolutionary paths, we incorporated the enzyme distributions into phylogenetic trees constructed via NCBI taxonomy identifiers [12]. Through a combinational analysis of the pathways in the phylogenetic trees, we have identified interesting distribution patterns of metabolic pathways that are linked with bacterial groups specific lifestyles, which may improve our understanding of the functions of energy reserves in bacteria. In addition, systematic analysis also gives us an overview of enzyme distributions, which could serve a as guide for further theoretical and experimental analyses of energy reserves in bacteria.

**Table 1.** Key enzymes and corresponding UniProt sequences used in this study for statistical modelling via HMMER package.

| Reference Species | Gene | Enzyme | Length | UniProt ID | Pfam Domain ID |
| --- | --- | --- | --- | --- | --- |
| *Escherichia coli* | *ppk1* | Polyphosphate kinase | 688 | E7QTB5<sup>#</sup> | PF02503, PF13090, PF17941, PF13089 |
| *Mycobacterium tuberculosis* | *ppk2* | Polyphosphate kinase 2 | 295 | O05877 | PF03976 |
| *Escherichia coli* | *ppx* | Ppx/GppA phosphatase | 513 | P0AFL6 | PF02541 |
| *Escherichia coli* | *glgC* | Glucose-1-phosphate adenylyltransferase | 431 | P0A6V1 | PF00483 |
| *Escherichia coli* | *glgA* | Glycogen synthase | 477 | P0A6U8 | PF08323, PF00534 |
| *Thermococcus kodakaraensis* | *glgB*<sup>*</sup> | Apha-1,4-glucan branching enzyme (GH57) | 675 | Q5JDJ7<sup>#</sup> | PF09210, PF03065, PF14520 |
| *Escherichia coli* | *glgB* | Alpha-1,4- glucan branching enzyme (GH13) | 728 | P07762<sup>#</sup> | PF02922, PF00128, PF02806 |
| *Mycobacterium tuberculosis* | *treS* | Trehalose synthase/amylase | 601 | P9WQ19<sup>!</sup> | PF00128, PF16657 |
| *Mycobacterium tuberculosis* | *pep2* | Maltokinase | 455 | Q7DAF6<sup>!</sup> | PF18085 |
| *Mycobacterium tuberculosis* | *glgE* | Alpha-1,4-glucan: maltose-1-phosphate Maltosyltransferase | 701 | P9WQ17<sup>!</sup> | PF00128, PF11896 |
| *Mycobacterium tuberculosis* | *Rv3032* | Glycogen synthase | 414 | P9WMY9 | PF13439, PF00534 |
| *Cupriavidus necator* | *phaA* | Acetyl-CoA acetyltransferase | 246 | P14611 | PF02803, PF00108 |
| *Cupriavidus necator* | *phaB* | Acetoacetyl-CoA reductase | 393 | P14697 | PF00106 |
| *Allochromatium vinosum* | *phaC* Group 1<sup>&</sup> | Poly(3-hydroxyalkanoate) polymerase subunit C | 355 | P45370 | PF00561 |
| *Pseudomonas aeruginosa* | *phaC* Group 2<sup>&</sup> | Class II poly(R)-hydroxyalkanoic acid synthase | 559 | Q51513 | PF07167 |
| *Escherichia coli* | *fabG* | 3-Oxoacyl-[acyl-carrier-protein] reductase | 244 | P0AEK2 | PF13561 |
| *Pseudomonas aeruginosa* | *phaJ* | (R)-Enoyl-CoA hydratase/enoyl-CoA hydratase I | 156 | Q9LBK2 | PF01575 |
| *Escherichia coli* | *fabD* | Malonyl CoA-acyl carrier protein transacylase | 209 | P0AAI9 | PF00698 |
| *Clostridium kluyveri* | *sucD* | Succinic semialdehyde dehydrogenase | 453 | P38947 | PF00171 |
| *Clostridium kluyveri* | *4hbD* | NAD-dependent 4-hydroxybutyrate dehydrogenase | 371 | P38945 | PF00465 |
| *Clostridium kluyveri* | *orfZ* | 4-Hydroxybutyrate CoA-transferase | 437 | A0A1L5FD42 | PF02550, PF13336 |
| *Saccharomyces cerevisiae* | *PDAT*<sup>^</sup> | Phospholipid: diacylglycerol acyltransferase | 661 | P40345 | PF02450 |
| *Acinetobacter baylyi* | *wax-dgaT* | Wax Ester Synthase/Acyl Coenzyme A:<br>Diacylglycerol Acyltransferase | 458 | Q8GGG1 | PF06974, PF03007 |

\* Archaeal type of glycogen branching enzyme (encoded by *glgB* gene) belonging to GH57 class in CATH database. #Hidden Markov models of protein sequences with more than 2 non-redundant domains were constructed from scratch.<sup>1</sup>HMMs of TreS, Pep2, and GlgE were constructed from scratch for full sequence models due to the existence of very short PFAM domains (Malt\_amylase\_C and Mak\_N\_cap) or unknown function PFAM domain (DUF3416) in their sequences. &*phaC* Group 1 with PFAM domain Abhydrolase\_1 (PF00561) and Group 2 with PFAM domain PhaC\_N (PF07167) should not be confused with the four types of PHA synthases that are classified based on primary sequences, substrate specificity, and subunit composition. ^TAG synthesis enzyme *PDAT* dominantly present in eukaryotes and only 42 out of 8282 bacterial proteomes show single or double homologs of the enzyme.

## Materials and Methods

### Proteomes and enzymes collection

Bacterial proteomes were downloaded from UniProt database by using two keywords, *Bacteria* and *Reference Proteomes*, as filters [11]. A total of 8282 bacterial proteomes were collected, reflecting the state of knowledge as at 2018. Of these, 68 proteomes were removed due to outdated taxonomy identifiers that cannot be identified in NCBI taxonomy database when constructing phylogenetic trees [12]. A complete list of all the 8282 bacteria with UniProt proteome identifiers, NCBI taxonomy identifiers, bacterial names, proteome sizes, and distribution patterns of key enzymes is available in the **Supplementary Table S1**. For each of the five major energy reserves, only key enzymes were considered. For the synthesis of WE and TAG, the bifunctional enzyme wax ester synthase/acyl-CoA:diacylglycerol acyltransferase (WS/DGAT) was studied due to its pivotal role in these processes [10]. In addition, the enzyme phospholipid: diacylglycerol acyltransferase (PDAT) that catalyses the acyl-CoA-independent formation of triacylglycerol in yeast and plants were also screened in bacterial proteomes [13]. For metabolism of polyP, the key enzyme polyphosphate kinase (PPK1) for synthesis and two degradation enzymes, intracellular polyphosphate kinase 2 (PPK2) and extracellular Ppx/GppA phosphatase (PPX), were included [2]. For glycogen metabolism, two synthesis pathways were considered. The first one involves glucose-1-phosphate adenylyltransferase (GlgC), glycogen synthase (GlgA), and glycogen branching enzyme (GlgB) [4]. The second pathway includes trehalose synthase/amylase (TreS), maltokinase (Pep2), α-1,4-glucan: maltose-1-phosphate maltosyltransferase (GlgE), and glycogen branching enzyme (GlgB) [14]. Key enzyme Rv3032 for the elongation of α-1,4-glucan in another pathway relevant to glycogen metabolism and capsular glucan was included [14]. In addition, archaeal type GlgB belonging to the glycosyl hydrolase family 57 (GH57) was also investigated for comparative analysis due to its importance. As for PHAs, major enzymes responsible for synthesis such as Acetyl-CoA acetyltransferase (PhaA), acetoacetyl-CoA reductase (PhaB) and poly(3-hydroxyalkanoate) polymerase subunit C (PhaC) have been identified [15, 16]. Based on Pfam analysis, PhaC could be further divided into two groups, PhaC Group 1 with PFAM domain Abhydrolase_1 (PF00561) and Group 2 with PFAM domain PhaC_N (PF07167). In addition, other enzymes in the PHAs synthesis pathway were also studied, which are 3-oxoacyl-[acyl-carrier-protein] reductase (FabG), (R)-Enoyl-CoA hydratase/enoyl-CoA hydratase I (PhaJ), malonyl CoA-acyl carrier protein transacylase (FabD), succinic semialdehyde dehydrogenase (SucD), NAD-dependent 4-hydroxybutyrate dehydrogenase (4HbD), and 4-hydroxybutyrate CoA-transferase (OrfZ) [17]. Together with poly(3-hydroxyalkanoate) polymerase subunit C, these enzymes are able to synthesize PHAs in bacteria. All the enzymes are screened for presence and absence in collected bacterial proteomes. For details of these enzymes, please refer to **Table 1**.

### De novo construction of HMMs

All selected seed proteins were used for constructing statistical sequence models based on HMMs via HMMER package [18]. After obtaining sequences for all seed proteins from UniProt database, remote BLAST was performed to collect homologous sequences for each seed protein from the NCBI non-redundant database of protein sequences [19]. Usearch was used to remove the homologous sequences with more than 98% similarity from the selected proteins [20]. The standalone command-line version of MUSCLE was used so the multiple-sequence alignments were created automatically [21]. Heads or tails of multiple sequence alignments tend to be more inconsistent [22]. Thus, all MSAs were manually edited to remove heads and tails by using JalView [23]. HMMER was selected for the construction of HMMs through hmmbuild command. Since HMMER only recognizes STOCKHOLM format, all MSAs results were converted from FASTA to STOCKHOLM format. In addition, another set of HMM models sourced directly from PFAM database based on the domain structures of seed proteins were collected. For seed proteins with more than two domains, de novo constructed HMM models were used for homologous screening, while those with two or less domains, established PFAM domains were used. For searching homologs in bacterial proteomes, routine procedures were performed by following HMMER User’s Guide <u>eddylab.org/software/hmmer3/3.1b2/Userguide.pdf</u>. Only those hits meeting the criteria of E-value less than 1e-10 and hit length greater than 60% of query domains will be considered. Results obtained from HMM screening can be found in **Supplementary Table S1**. In particular, the presence (copy numbers) or absence of a specific enzyme in a certain bacterial proteome is noted, together with corresponding E-values and target sequence lengths.

### Data visualization

Phylogenetic trees were first constructed based on NCBI taxonomy identifiers for all bacteria in this study via the commercial web server PhyloT https://phylot.biobyte.de/, and were then visualized through the online interactive Tree of Life (iTOL) server https://itol.embl.de/ [24]. Distribution patterns of enzymes and their combinations in terms of energy reserves were added to the trees through iTOL pre-defined tol_simple_bar template [24].

### Statistical analysis

Unpaired two-tailed Student’s *t*-test was used for statistical analysis through R package. Significant difference was defined as *P-value* < 0.05.

## Results

### Wax ester and triacylglycerol

The key enzyme that is involved in both WE and TAG synthesis in bacteria is WS/DGAT. Through the screening of HMM-based statistical models, 673 out of 8282 bacterial species harbour a single copy or multiple copies of WS/DGAT homologs, which are mainly present in phylum *Actinobacteria* and the super-phylum *Proteobacteria*. Only a few bacteria in groups such as FCB (also known as *Sphingobacteria*) and PVC (also known as *Planctobacteria*), *etc*. have WS/DGAT. No species belonging to phylum *Firmicutes* has WS/DGAT. As for the unclassified bacteria, although no WS/DGAT was identified, they will not be studied due to their uncertain classification at current stage. For details, please refer to **Figure 1**. By comparing the proteome sizes of bacteria species with or without WS/DGAT, we found that bacteria with WS/DGAT have average proteome size of 5294 proteins/proteome while those without WS/DGAT have average proteome size of 3117 proteins/proteome (*P-value* < 0.001).

**Figure 1.**
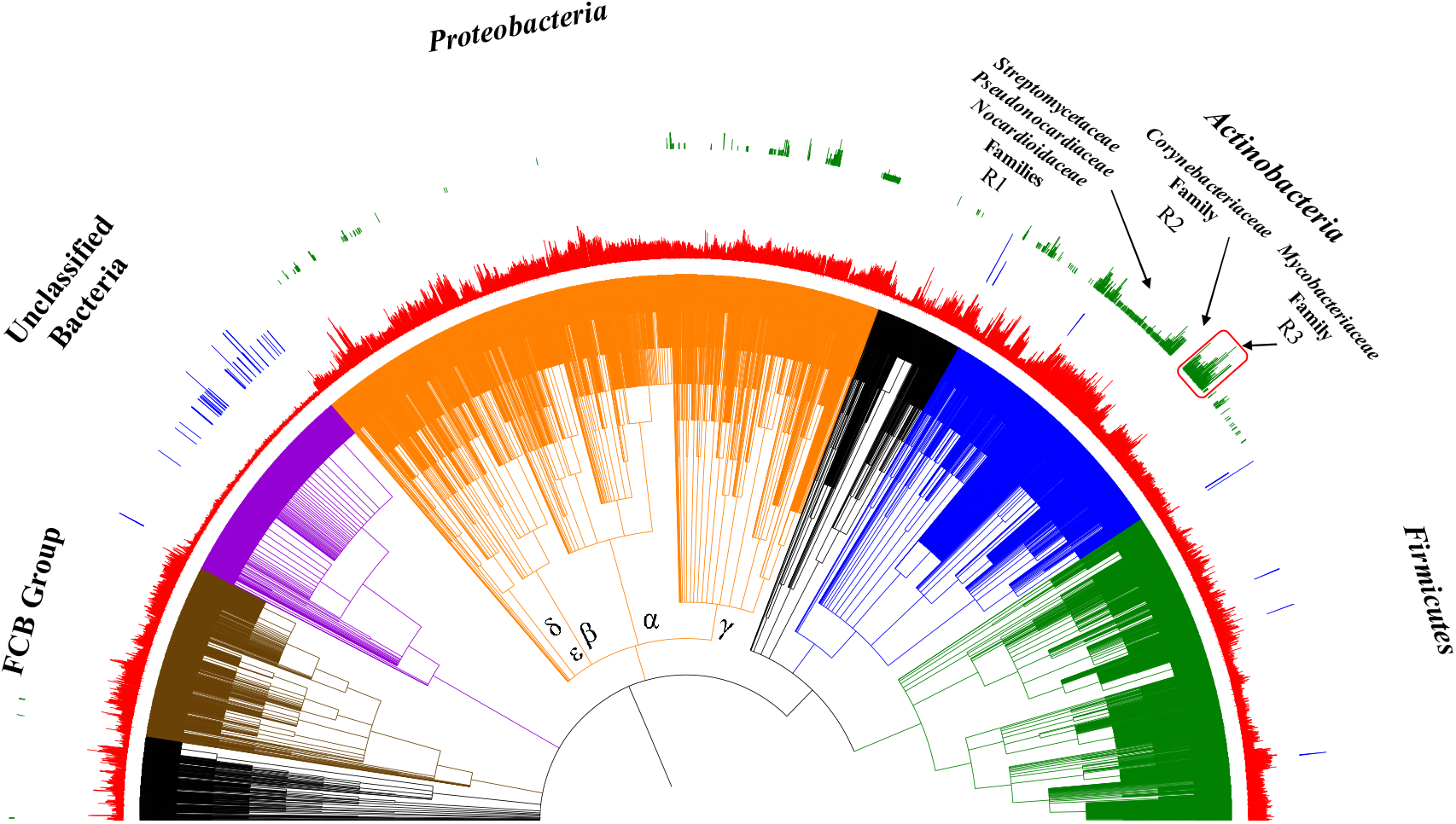
Analysis of the distribution patterns of key enzyme WS/DGAT (green circle) and PDAT (blue circle) along the evolutionary tree that is responsible for the synthesis of neutral lipids wax ester and triacylglycerol, together with bacterial proteome sizes (red circle). Most WS/DGATs were present in *Proteobacteria* (γ, δ, and ε subdivisions) and *Actinobacteria* phyla. Region 1 (R1) including three closely related families with most species harboring WS/DGATs were annotated, which included *Streptomycetaceae*, *Pseudonocardiaceae*, and *Nocardioidaceae*. Region R3 with high-copy WS/DGATs was highlighted with red rectangle, which exclusively fall into the *Mycobacteriaceae* family. Intriguingly, all selected 68 species in *Corynebacteriaceae* family (R2) that is closely related with *Mycobacteriaceae* family do not have any WS/DGAT. 46 PDAT homologs from 42 bacterial species were scarcely identified mainly in unclassified bacteria and *Terrabacteria* group (blue bar). Five groups of bacteria are highlighted, which are *Firmicutes* (green), *Actinobacteria* (blue), *Proteobacteria* (orange), Unclassified Bacteria (violet), and FCB group (brown).

Within the major phylum of *Proteobacteria*, WS/DGAT is not evenly distributed and γ-, δ-, and ε-*Proteobacteria* sub-divisions have more species harbouring WS/DGAT genes. In addition, two orders, *Rhodobacterales* (305 species) and *Enterobacterales* (168 species) that belong to α- and γ-*Proteobacteria* phylum, respectively, do not have any WS/DGAT except for one species *Plesiomonas shigelloides 302-73* (NCBI taxonomy ID 1315976). As for the phylum *Actinobacteria*, two WS/DGAT abundant regions (R1 and R3) and one WS/DGAT absence region (R2) in the phylogenetic tree are worth further exploration. R1 region includes three closely related families with most species harboring WS/DGATs, which includes *Streptomycetaceae*, *Pseudonocardiaceae*, and *Nocardioidaceae*. R3 includes only one family *Mycobacteriaceae* (115 species) in which bacteria have up to 10 homologs of WS/DGAT. R4 is the family *Corynebacteriaceae* (69 species) that only includes WS/DGAT-free bacteria.

As for phospholipid:diacylglycerol acyltransferase (PDAT), it is involved in TAG storage in yeasts and plants. Recently, PDAT activity was proven in *Streptomyces coelicolor* for the first time [25]. However, no bacterial homologs were known yet with similarity to the respective eukaryotic sequences [26]. Our systematic screening of *Saccharomyces cerevisiae* PDAT unexpectedly identified 46 proteins belonging to 42 out of 8282 bacterial species (**Supplementary Table 2**), 38 homologs belong to small genome-sized unclassified bacteria while 8 homologs were present in *Terrabacteria* group. Both E-values (>1E-10) and target sequence lengths (> 60% query sequence length) were also given to confirm the significance of the results. Thus, presence of eukaryotic PDAT in bacteria is theoretically true, though functionality of the enzymes requires further experimental validation. Although most of the sequences were annotated as uncharacterized proteins in the UniProt database, some were described as acyltransferases. Interestingly, the PDAT homolog in *Clostridium butyricum* E4 str. BoNT E BL5262 was thought to be a putative prophage LambdaBa01 acyltransferase, which indicated that the enzyme could jump around among species via horizontal gene transfer.

### Polyhydroxyalkanoates

Although TAG and WE are a more common storage lipid in several groups of bacteria, the majority of all bacteria store PHA rather than TAG or WE [27]. Three major enzymes (PhaA, PhaB and PhaC) involved in the synthesis of PHAs, forming the classical synthesis pathway and two enzymes (intracellular PhaZ and extracellular PhaZ) involved in the utilization of PHAs in bacteria. In this study, we mainly focused on the distribution patterns of PHA synthesis pathway PhaABC. PhaC has been further divided into four classes depending on substrate specificities and subunit compositions, represented by *Cupriavidus necator* (class I), *Pseudomonas aeruginosa* (class II), *Allochromatium vinosum* (class III), and *Bacillus megaterium* (class IV) [28]. However, based on Pfam analysis, the four classes of PhaC could be included into two groups, PhaC Group 1 with PFAM domain Abhydrolase_1 (PF00561) and Group 2 with PFAM domain PhaC_N (PF07167), which were also thoroughly investigated for their distribution patterns. However, PhaE and PhaR as PhaC subunits were not be considered in this study. On the other hand, there are another four confirmed pathways for PHAs synthesis, which are 1) FabG, 2) PhaJ, 3) FabD, 4) SucD, 4HbD and OrfZ [17]. All the enzymes are screened for presence and absence in collected bacterial proteomes (**Supplementary Table 1**).

Preliminary analysis showed that 3166 bacterial species with average proteome size of 3772 proteins/proteome have PhaABC pathway while 1036 bacterial species with average proteome size of 1059 proteins/proteome do not have the pathway (*P-value*<0.001). In general, PHA synthesis was widely distributed across bacterial species. It is interesting to notice that Group II PHA synthase is dominantly present in *Proteobacteria* Phylum while Group I PHA synthase is abundantly present in *Actinobacteria* Phylum. In addition, three representative regions R1, R2, and R3 with loss of the classical PHA synthesis pathway were highlighted, that is, *Bacteroidia* Class, Epsilon-*Proteobacteria* Subdivision Class, and *Tenericutes* Phylum. In these regions, two bacterial species, *Lentimicrobium saccharophilum* and *Helicobacter* sp. 13S00477-4 actually harboured the complete PhaABC pathway. Distribution patterns of the other four PHA synthesis pathways mentioned above were not visualized along phylogenetic tree since they were distributed with no featured patterns (**Supplementary Table 1**). Please refer to **Figure 2** for the distribution patterns of the studied enzymes and the pathway.

**Figure 2.**
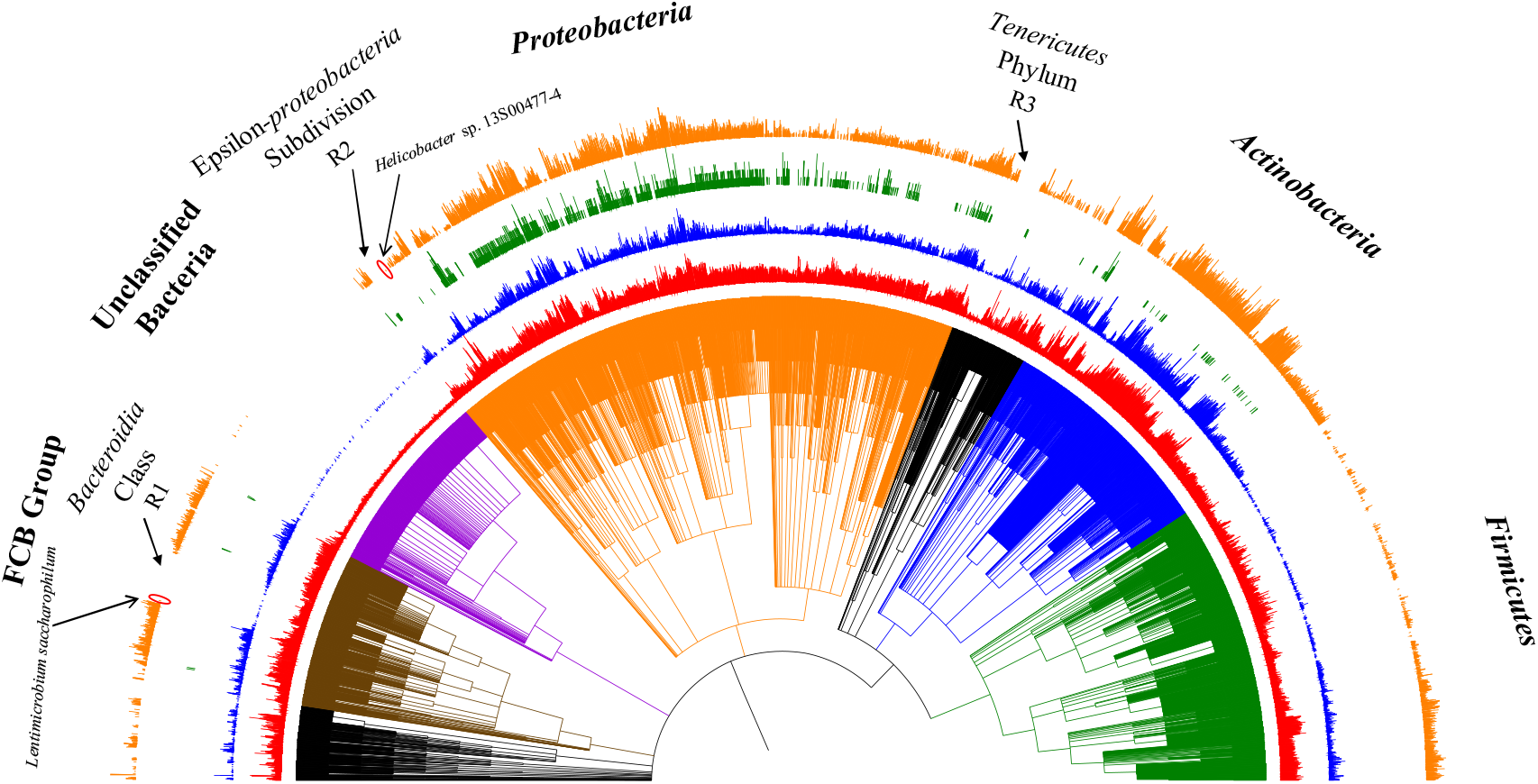
Distribution patterns of the classical PHA synthesis pathway PhaABC (orange circle) and the two groups of PHA synthases, Group I with Abhydrolase_1 domain (blue circle) and Group II with PhaC_N domain (green circle). Bacterial proteome sizes were presented in red bar. Three representative regions (R1, R2, and R3) with the loss of PHA synthesis pathway are highlighted, which belong to *Bacteroidia* Class, Epsilon-*Proteobacteria* Subdivision, and *Tenericutes* Class. In addition, Group II PHA synthase showed a skewed distribution pattern with dominant existence in *Proteobacteria* Phylum while Unclassified Bacteria Group and *Firmicutes* Phylum do not have any homolog of the enzyme.

### Polyphosphate

Three key enzymes PPK1, PPK2, and PPX are related with polyP metabolism. PPK1 is responsible for polyP synthesis. In this study, a total of 5273 bacterial species have PPK1. PPK2 and PPX are used for intracellular and extracellular polyP degradation, respectively. 2584 bacterial species have PPK1, PPK2 and PPX enzymes while 2211 bacteria species do not have any of the three enzymes. The average proteome sizes of the two groups of bacteria are 4571 proteins/proteome and 1615 proteins/proteome, respectively, which are significantly different (*P-value* < 0.001). In our analysis, we independently reviewed the distribution patterns of the three enzymes along phylogenetic trees and the result is displayed in **Figure 3**. The three enzymes are widely distributed across bacterial species, which reflects the essentiality of the polymer in bacterial physiology. In addition, comparison shows that *Firmicutes* phylum seems to favour PPX more than PPK2 for polyP degradation. In addition, although it was observed that several regions had missing synthesis enzyme or degradation enzyme, only unclassified bacteria and *Mollicutes* class (94 bacterial species) showed apparent lack of the three polyP metabolism enzymes.

**Figure 3.**
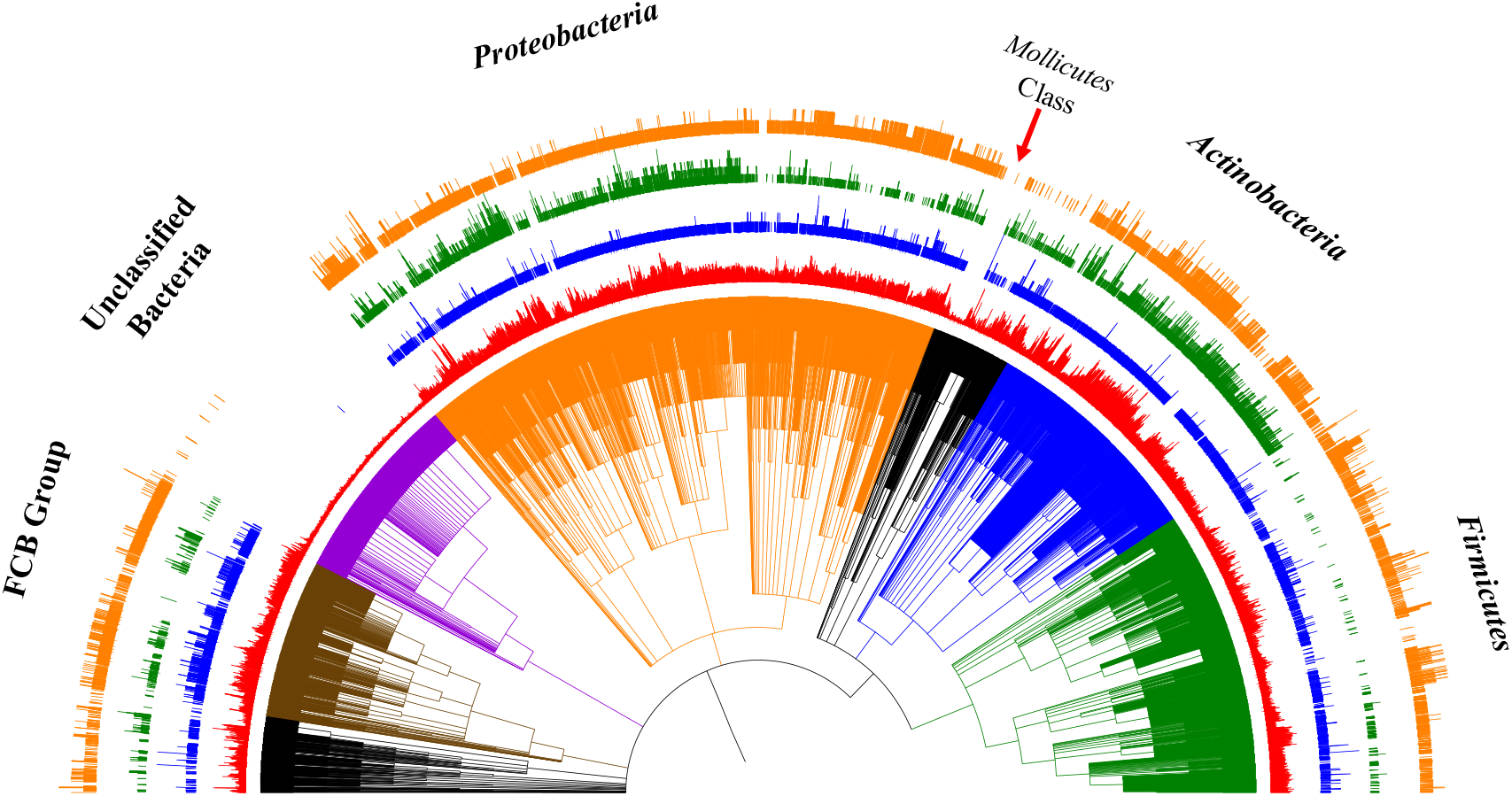
Distribution patterns of key enzyme polyphosphate kinase PPK1 (blue circle), polyphosphate kinase 2 PPK2 (green circle), and exopolyphosphatase PPX (orange circle) along the evolutionary pathway that is responsible for main synthesis and degradation pathways of polyphosphate in bacteria. Polyphosphate metabolism is widely distributed in bacteria. Only unclassified bacteria and the class of Mollicutes belonging to phylum Tenericutes have apparent loss of polyphosphate metabolism.

### Glycogen

Glycogen metabolism in bacteria has multiple pathways, which include the classical pathway (GlgC, GlgA, GlgB, GlgP and GlgX) [4], trehalose pathway (TreS, Pep2, GlgE, and GlgB), and the novel Rv3032 pathway [29]. Rv3032 is an alternative enzyme for the elongation of α-1,4-glucan, which was compared for distribution patterns with GlgA. We also focused on the two glycogen synthesis pathways and compared their distribution patterns. In addition, there are two types of glycogen branching enzymes. One is the common bacterial GlgB, belonging to GH13 in CATH database, and the other one is known as archaeal GlgB, belonging to GH57 in CATH database [30]. We also looked into their distribution patterns in bacteria since GlgB is essential in determining the branched structure of glycogen. Our study showed that 3137 bacteria have the classical synthesis pathway (GlgC, GlgA, and GlgB) and their average proteome size is 3966 proteins/proteome while only 510 bacterial species (average proteome size of 1120 proteins/proteome) do not have these enzymes (*P-value*<0.001). Comparison of the two synthesis pathways confirmed that classical synthesis pathways are widely distributed across species. Random loss of the classical pathway can be inferred from **Figure 4**. In contrast, trehalose pathway is tightly associated with *Actinobacteria* phylum. It was also shown that Rv3032 was actually more widespread than GlgA. As for the two GlgBs, GH13 GlgB is widely distributed in 4500 bacterial species with a trend of random loss, while GH57 GlgB is identified in only 785 bacterial species that are mainly fall into *Terrabacteria* Group and PVC group, *etc*.

**Figure 4.**
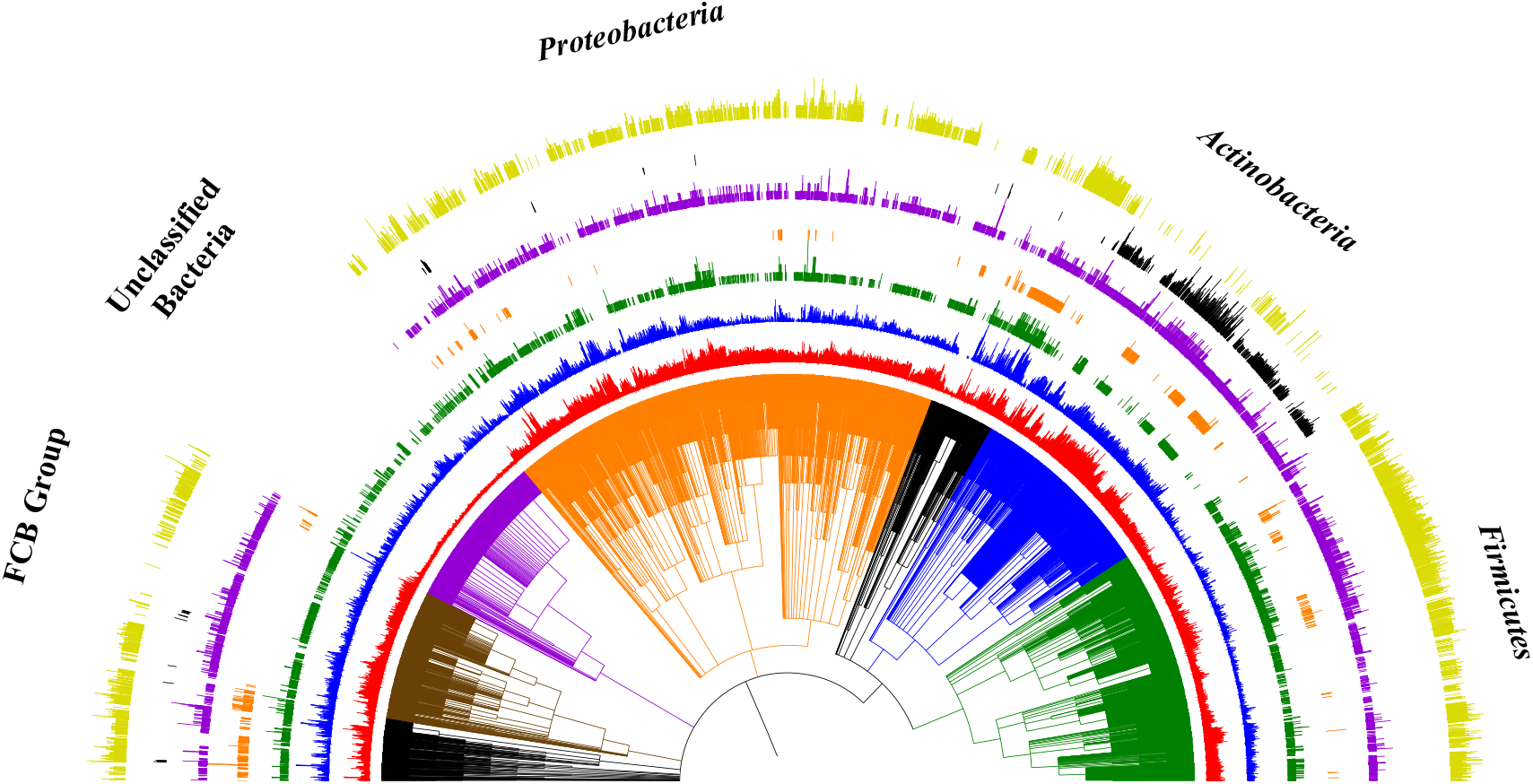
Distribution patterns of two glycogen synthesis pathways and four key enzymes. Full classical glycogen synthesis pathway (yellow circle) includes GlgC, GlgA, and GlgB, which distributes widely in bacteria except for *Actinobacteria* phylum, FCB group and unclassified bacteria group. The other one is trehalose-based glycogen synthesis pathway including TreS, Pep2, GlgE, GlgB (black circle), which is mainly restricted to the *Actinobacteria* phylum. As for the four key enzymes, glycogen synthases (Rv3032 in blue circle and GlgA in green circle) and glycogen branching enzymes (GH57 GlgB in orange circle and GH13 GlgB in violet circle) were analyzed along the NCBI taxonomy tree. As for Rv3032, its distribution is more widely present than GlgA in almost all bacteria. Distributions of the two GlgB enzymes show that GH57 GlgB is mainly present in *Terrabacteria*. In addition, GH57 GlgB is also identified in PVC group, *Spirochaetes*, *Acidobacteria*, *Fusobacteria*, *Thermotogae*, *Nitrospirae*, *Aquificae*, *Synergistetes*, *Elusimicrobia*, *Nitrospinae/Tectomicrobia* group, *Thermodesulfobacteria*, *Rhodothermaeota*, and *Dictyoglomi*.

## Discussion

From an evolutionary point of view, if an organism can obtain compounds from other sources, it would tend to discard the corresponding biosynthetic pathways [31]. For example, strictly intracellular bacteria such as *Rickettsia* species, *Mycoplasma* species, and *Buchnera*, *etc.* have extensively reduced genome sizes and common pathways relevant to energy metabolism have been eliminated [31]. Although a common belief is that organism should evolve toward high complexity, recent analysis reported that reduction and simplification could be the dominant mode of evolution while increasing complexity is just a transitional stage according to the neutral genetic loss and streamlining hypothesis [32]. Independent analyses of the distribution patterns of the five energy reserves in bacteria found a consistent and statistically significant correlation between energy reserve loss and reduced proteome size. Previous studies also reported this correlation in terms of glycogen and polyP metabolism in bacteria [2, 4]. In this study, we extended the conclusion by adding the neutral lipid reserves of WE, TAG, and PHA. It was also confirmed that bacteria losing energy reserve metabolism capacity tended to have a niche-dependent or host-dependent lifestyle [2, 4, 6]. Thus, by looking into bacterial energy reserve metabolism, we could obtain preliminary views in terms of bacterial lifestyles, though other evidence is required to verify this hypothesis. It is worth mentioning that proteomes that we used in this study were derived from translated coding genes, not based on condition-specific expression of proteins [11]. Thus, there is no bias in protein coverage or gene expression level. From this point of view, this equivalent to bacterial genome analysis, except for that HMM models perform much better for protein sequences than DNA sequences in terms of remote homolog identification.

WS/DGAT is a bifunctional enzyme and key to the biosynthesis of WE and TAG in bacteria. It was previously thought that WE and TAG are very uncommon lipid storage compounds in bacteria when compared with plants and animals, until this novel enzyme was identified [10]. From our analysis, it could be seen that many bacteria belonging to both Gram-positive and Gram-negative categories have the potential to synthesize WE and TAG. However, studies about WS/DGAT are mainly restricted to *Mycobacteria* genus (*Actinobacteria* phylum) and *Acinetobacter* genus (γ-Proteobacteria phylum) due to their clinical significance and potentially industrial use. WS/DGAT genes in the phylum *Actinobacteria* tend to have more paralogs than other phyla, especially for the bacteria in the R1 and R3 regions, which include *Mycobacteriaceae*, *Dietziaceae*, *Gordoniaceae*, *Nocardiaceae*, *Tsukamurellaceae*, *Williamsiaceae*, *Nocardioidaceae* and *Pseudonocardiales*. On the other hand, no WS/DGAT is found in the family of *Corynebacteriaceae* (R2 region), although *Corynebacteriaceae* is closely related with *Mycobacteriacea* [33]. In addition, bacteria in phylum *Firmicutes* do not have any WS/DGAT enzymes. Screening of Phospholipid:diacylglycerol acyltransferase (PDAT), an enzyme that catalyses the acyl-CoA-independent formation of triacylglycerol in yeast and plants, found 46 homologs in bacteria [13]. This contradicts with previous thoughts that eukaryotic PDAT is exclusively present in higher organisms with no homologs in prokaryotic genomes [27].

The family *Corynebacteriaceae* contains the genera *Corynebacterium* and monospecific genus *Turicella* [34]. *Mycobacterium tuberculosis* is the dominant species in *Mycobacteriaceae* (97 species). Mycolic acid (MA), with wax ester as the oxidized form of MA in *Mycobacterium tuberculosis*, is present in the cell wall and plays essential roles in host invasion, environmental persistence, and also drug resistance [35]. In addition, *Mycobacterium tuberculosis* also relies on wax ester for dormancy, though specific functions of WE in *M. tuberculosis* require further investigation. Thus, abundance of WS/DGAT in *Mycobacteriacea* has selective advantages in evolution. Considering the abundance of wax ester and its slow degradation, it could also contribute to the long-term survival (more than 360 days) of *M. tuberculosis* in environment [6]. On the other hand, *Corynebacterium* does not rely on oxidized mycolic acid while *Turicella* does not have mycolic acid at all [36, 37]. Thus, there is no need for them to be equipped with the WS/DGAT enzyme. However, how *Mycobacteriaceae* gains WS/DGAT, or *Corynebacteriaceae* loses it, is not clear and needs more investigation. As for *Firmicutes*, it is the low G+C counterpart of the high G+C *Actinobacteria*. Most of its species can form endospores and are resistant to extreme environmental conditions such as desiccation, temperature fluctuation, and nutrient deprivation, *etc.* [38]. Thus, they may not need compounds such as WE or TAG for storing energy and dealing with harsh external conditions. How G+C content in the two phyla may impact on the gain or loss of wax ester metabolism is currently not known.

PHAs are a group of compounds that include but are not limited to components such as polyhydroxybutyrate (PHB) and polyhydroxyvalerate (PHV), *etc*., among which PHB is the most common and most prominent member in bacteria [39, 40]. Currently, there are eight pathways responsible for the synthesis of PHB in bacteria [17]. The complexity of the metabolic pathways in bacteria is worthy of a separate and complete investigation and is unable to be fully addressed in this study. Here, we only focused on the classical pathway that mainly involves PhaA, PhaB and PhaC [39]. Its phylogenetic analysis revealed that PHA synthesis is widely distributed across bacterial species, which is consistent with its role as a dominant neutral lipid reserve in bacteria. As discussed above, PHA synthase is further divided into four classes. However, HMM analysis of representative sequences revealed that differences of the four-class enzymes at domain level only involves Abhydrolase_1 (PF00561) and PhaC_N (PF07167) domains. Further exploration of all bacterial PhaC domain structures shows higher heterogeneity, indicating that a more complex classification system for this enzyme should be introduced (unpublished data) and will be investigated in future study.

PolyP is known to be ubiquitous in different life domains and claimed to be present in all types of cells in Nature due to its essential roles as energy and phosphate sources [2]. Although a number of enzymes are directly linked with polyP metabolism, we only focused on PPK1, PPK2, and PPX in this study because they are most essential enzymes to polyP metabolism. **Figure 3** gives an overview of the distribution patterns of the three enzymes. Although 2212 bacterial species across the phylogenetic tree lack of all three enzymes, an apparent gap was only evident in the phylum *Tenericutes* and was further confirmed to be *Mollicutes*. A previous analysis of 944 bacterial proteomes showed that bacteria which have completely lost the polyP metabolic pathways (PPK1, PPK2, PAP, SurE, PPX, PpnK and PpgK) are heavily host-dependent and tend to adopt obligate intracellular or symbiotic lifestyles [2]. Consistently, *Mollicutes* is a group of parasitic bacteria that have evolved from a common *Firmicutes* ancestor through reductive evolution [41]. From here, we could infer that not only loss of complete metabolism pathway, but also even loss of key enzymes for energy reserve metabolism could give a hint at bacterial lifestyle.

For glycogen metabolism, we compared two synthesis pathways, the classical pathway (GlgC, GlgA and GlgB) and the newly identified trehalose-related pathway (TreS, Pep2, GlgE and GlgB) [4, 14]. Although initial analysis via BLAST search showed in 2011 that GlgE pathway is represented in 14□% of sequenced genomes from diverse bacteria, our studies showed that, when searching for the complete GlgE pathway by including another three enzymes, it is dominantly restricted to *Actinobacteria* phylum while the classical pathway is widely present in the phylogenetic tree as seen in **Figure 4** [14]. In addition, the two types of GlgBs also showed interesting distribution patterns. Although GH13 GlgB is widely identified in 54.33% bacteria, GH57 GlgB is only present in 9.48% bacteria with skewed distribution in groups such as the *Terrabacteria* phylum and PVC group, *etc*. Another study of 427 archaea proteomes found that the 11 archaea have GH13 GlgB, while 18 archaea have GH57 GlgB [29]. Thus, the two GlgBs are rarely present in archaea and mainly exist in bacteria. However, why trehalose-related glycogen metabolism pathway is markedly associated with *Actinobacteria* phylum still needs more experimental exploration.

## Conclusions

Distribution patterns of key enzymes and their combined pathways in bacteria provided a comprehensive view of how energy reserves are incorporated and lost during evolutionary processes. In general, polyP, PHA, and glycogen are widely distributed across bacterial species as energy storage compounds. The other two neutral lipids investigated in this study are comparatively minor energy reserves in bacteria and mainly found in the super phylum *Proteobacteria* and phylum *Actinobacteria*. Within the group, more bacteria have the capacity to accumulate WE and TAG due to the abundance of WS/DGAT homolog. Comparatively. polyP acts as a transient energy reserve while neutral lipids are a more sustainable energy provider [4, 42]. Thus, neutral lipids could be major players for bacterial persistence under harsh conditions such terrestrial and aquatic environments. As for glycogen, its ability to enhance bacterial environmental viability is still controversial. Its widespread distribution in bacteria indicates that its metabolism is tightly linked with bacterial essential activities. In sum, through this study, we obtained a much clearer picture about how key enzymes responsible for the metabolism of energy reserves are distributed in bacteria. Further investigation via incorporating bacterial physiology and lifestyle could supply additional explanations to illustrate the distribution patterns, although experimental evidence is indispensable to confirm the computational analysis.

## Acknowledgements

We thank Professor Ximiao He from Huazhong University of Science and Technology and the anonymous reviewers for genuine comments and suggestions, which greatly improves the quality of this study. The work was supported by Startup Foundation for Excellent Researchers at Xuzhou Medical University [D2016007], Innovative and Entrepreneurial Talent Scheme of Jiangsu Province (2017), and Nature and Science Foundation of Jiangsu Province [BK20180997].

## Author Contributions

LW conceived the core idea of this study. LW, MJW, PX, JY, YX, QL, and YH did all data collection, data visualization, and statistical analysis. LW and MJW created and edited the manuscript.

## Declaration of Conflicting Interests

The authors declare that there is no conflict of interest.

