## Supplementary Table 2 for "Systematic Analysis of Metabolic Pathway Distributions of Bacterial Energy Reserves"

**Supplementary Table 2** Homologous sequences of the fungal enzyme Phospholipid: diacylglycerol acyltransferase (PDAT) in bacteria based on HMM screening.

| **Proteome ID** | **Taxonomy ID** | **Organism** | **Protein Size** | **UniProt ID** | **Protein** |
| --- | --- | --- | --- | --- | --- |
| UP000003081 | 632245 | *Clostridium butyricum* E4 str. BoNT E BL5262 | 4245 | C4IEU2 | Putative prophage LambdaBa01 Acyltransferase |
| UP000005039 | 1095750 | *Lachnoanaerobaculum saburreum* F0468 | 3005 | I0R882 | von Willebrand factor type A domain protein |
| UP000006321 | 1206781 | *Paenibacillus alvei* DSM 29 | 6508 | K4ZD00 | Putative regulator of chromosome condensation RCC1 |
| UP000029539 | 1284708 | *Tissierellia bacterium* S7-1-4 | 1482 | A0A095ZF37 | Uncharacterized protein |
| UP000033988 | 1618902 | *Parcubacteria* group bacterium GW2011_GWC1_40_11 | 750 | A0A0G0QHI8  A0A0G0QQQ9 | Acetyltransferases and hydrolases with the alpha/beta hydrolase fold protein  Uncharacterized protein |
| UP000034191 | 1618894 | *Parcubacteria* group bacterium GW2011_GWC1_36_108 | 1009 | A0A0G0D8T9 | Acetyltransferases and hydrolases with the alpha/beta hydrolase fold protein |
| UP000034201 | 1618608 | Candidatus *Adlerbacteria* bacterium GW2011_GWC1_50_9 | 910 | A0A0G1YWE1 | Uncharacterized protein |
| UP000034272 | 1618793 | *Parcubacteria* group bacterium GW2011_GWA1_47_8 | 891 | A0A0G1TK15 | Cell surface protein |
| UP000034481 | 1618863 | *Parcubacteria* group bacterium GW2011_GWB1_37_13 | 506 | A0A0G0GD58 | YD repeat-containing protein |
| UP000034535 | 1618515 | *Microgenomates* group bacterium GW2011_GWC1_39_7 | 635 | A0A0G0RUX0 | Uncharacterized protein |
| UP000034605 | 1618715 | Candidatus *Moranbacteria* bacterium GW2011_GWF1_34_10 | 1233 | A0A0G0DS51 | Uncharacterized protein |
| UP000034654 | 1618915 | *Parcubacteria* group bacterium GW2011_GWC1_45_14 | 954 | A0A0G1PFT0  A0A0G1PA66 | Cell surface protein，Acetyltransferases and hydrolases with the alpha/beta hydrolase fold protein |
| UP000034759 | 1618892 | *Parcubacteria* group bacterium GW2011_GWC1_35_21 | 629 | A0A0G0CF87 | Acetyltransferases and hydrolases with the alpha/beta hydrolase fold protein |
| UP000034801 | 1618803 | *Parcubacteria* group bacterium GW2011_GWA1_56_13 | 759 | A0A0G1YJ47 | Uncharacterized protein |
| UP000034839 | 1618721 | Candidatus *Moranbacteria* bacterium GW2011_GWF2_35_39 | 1019 | A0A0G0EWU6 | Uncharacterized protein |
| UP000034889 | 1618657 | Candidatus *Giovannonibacteria* bacterium GW2011_GWC2_44_8 | 526 | A0A0G1M896 | Uncharacterized protein |
| UP000034945 | 1618707 | Candidatus *Moranbacteria* bacterium GW2011_GWE1_35_17 | 1299 | A0A0G0BUK5  A0A0G0DZA7 | Uncharacterized protein，Uncharacterized protein |
| UP000036787 | 1659198 | *Parcubacteria* bacterium C7867-004 | 851 | A0A0L0LA76 | Lecithin-cholesterol acyltransferase |
| UP000037457 | 1659200 | *Parcubacteria* bacterium C7867-006 | 685 | A0A0L0LCW1 | Lecithin-cholesterol acyltransferase |
| UP000176187 | 1801774 | Candidatus *Nomurabacteria* bacterium RIFCSPLOWO2_01_FULL_41_12 | 641 | A0A1F6WX61 | Uncharacterized protein |
| UP000176558 | 1802776 | Candidatus *Zambryskibacteria* bacterium RIFCSPLOWO2_12_FULL_39_23 | 601 | A0A1G2UTY5 | Uncharacterized protein |
| UP000176868 | 1802782 | Candidatus *Zambryskibacteria* bacterium RIFOXYD2_FULL_43_10 | 423 | A0A1G2V618 | Uncharacterized protein |
| UP000177235 | 1817845 | Candidatus *Doudnabacteria* bacterium RIFCSPLOWO2_02_FULL_48_13 | 904 | A0A1F5QCE4 | Uncharacterized protein |
| UP000177325 | 1798525 | Candidatus *Kaiserbacteria* bacterium RIFCSPLOWO2_12_FULL_45_26 | 961 | A0A1F6FFX4 | Uncharacterized protein |
| UP000177370 | 1801739 | Candidatus *Nomurabacteria* bacterium RIFCSPHIGHO2_01_FULL_40_24b | 997 | A0A1F6V7S2  A0A1F6V5Q9 | Uncharacterized protein  Uncharacterized protein |
| UP000177480 | 1802114 | Candidatus *Ryanbacteria* bacterium RIFCSPHIGHO2_01_FULL_45_22 | 1056 | A0A1G2FXN8 | Uncharacterized protein |
| UP000177682 | 1817838 | Candidatus *Doudnabacteria* bacterium RIFCSPHIGHO2_12_FULL_48_16 | 958 | A0A1F5PK07 | Uncharacterized protein |
| UP000177720 | 1798361 | Candidatus *Giovannonibacteria* bacterium RIFCSPLOWO2_12_43_8 | 557 | A0A1F5Y1H1 | Uncharacterized protein |
| UP000177777 | 1801754 | Candidatus *Nomurabacteria* bacterium RIFCSPHIGHO2_02_FULL_41_18 | 625 | A0A1F6W7L6 | Uncharacterized protein |
| UP000178042 | 1798490 | Candidatus *Kaiserbacteria* bacterium RIFCSPHIGHO2_02_FULL_49_16 | 910 | A0A1F6DD58 | Uncharacterized protein |
| UP000178114 | 1798351 | Candidatus *Giovannonibacteria* bacterium RIFCSPLOWO2_01_FULL_45_34 | 832 | A0A1F5X003 | Uncharacterized protein |
| UP000178184 | 1801764 | Candidatus *Nomurabacteria* bacterium RIFCSPLOWO2_01_FULL_33_17 | 565 | A0A1F6WP55 | Uncharacterized protein |
| UP000178367 | 1797994 | Candidatus *Falkowbacteria* bacterium RIFOXYA2_FULL_47_19 | 1531 | A0A1F5SMT8 | Uncharacterized protein |
| UP000178815 | 1798481 | Candidatus *Kaiserbacteria* bacterium RIFCSPHIGHO2_01_FULL_53_31 | 688 | A0A1F6CHP0 | Uncharacterized protein |
| UP000179230 | 1798532 | Candidatus *Kaiserbacteria* bacterium RIFOXYD1_FULL_42_15 | 678 | A0A1F6FPL9 | Uncharacterized protein |
| UP000187591 | 1739304 | *Anaerosphaera* sp. HMSC064C01 | 1440 | A0A1E9AKJ2  A0A1E9AFZ7 | Uncharacterized protein  Uncharacterized protein |
| UP000192674 | 2030 | *Kibdelosporangium* aridum | 11140 | A0A1W2FKP0 | Lecithin:cholesterol acyltransferase |
| UP000198313 | 2005459 | *Tolypothrix* sp. NIES-4075 | 7285 | A0A218QRZ4  A0A218QUM7 | Uncharacterized protein  Uncharacterized protein |
| UP000216537 | 1970248 | *Parcubacteria* group bacterium 21-54-25 | 837 | A0A257THS5 | Uncharacterized protein |
| UP000216926 | 1970249 | *Parcubacteria* group bacterium 21-58-10 | 758 | A0A257RQA4 | Uncharacterized protein |
| UP000218514 | 2005458 | *Nostoc* sp. NIES-4103 | 6898 | A0A1Z4RYC0 | Uncharacterized protein |
| UP000228524 | 1974675 | Candidatus *Moranbacteria* bacterium CG23_combo_of_CG06-09_8_20_14_all_39_10 | 921 | A0A2G9Z1R4 | Uncharacterized protein |
